# Phosphoinositide turnover through PLCγ regulates Draper-dependent engulfment in glia

**DOI:** 10.64898/2026.06.04.729572

**Authors:** Freya Storer, Eilish Mackinnon, Hannah Lucas-Clarke, Daniel Maddison, Leonardo Amadio, Thejaswini Susobhanan, Bilal R. Malik, Owen M. Peters, Gaynor A. Smith

## Abstract

Glial engulfment of degenerating neuronal material is essential for nervous system development, maintenance and repair. Genome-wide association studies have identified protective variants in the phosphoinositide-metabolising enzyme PLCG2 that modify Alzheimer’s disease risk, but how PLCG2-dependent phosphoinositide signalling regulates glial engulfment remains unclear. Using *Drosophila*, we investigated the role of *small wing (sl)*, the fly orthologue of human PLCG2, in glial responses to axonal injury and amyloid pathology. Glial knockdown of *sl* altered immune-associated transcriptional pathways and significantly delayed clearance of degenerating olfactory receptor neuron axons following axotomy. Loss of *sl* disrupted injury-induced phosphoinositide remodelling, resulting in elevated basal PIP2 levels and impaired post-injury accumulation of PIP3. Similar defects were observed following knockdown of the engulfment receptor Draper, placing phosphoinositide turnover downstream of Draper signalling. Simultaneous *Pten* knockdown restored phosphoinositide signalling and rescued delayed neuronal clearance in *sl*-deficient glia. Loss of *sl* also prevented injury-induced Draper upregulation and disrupted glial calcium signalling responses to axonal injury. In a model of Aβ42 accumulation, *sl* knockdown altered brain PIP2/PIP3 balance and improved survival independently of amyloid burden. Together, these findings identify PLCγ-dependent phosphoinositide turnover as a conserved regulator of Draper-mediated glial engulfment and provide mechanistic insight into how PLCG2 influences glial function and neurodegenerative disease risk.

**Graphical Abstract:** 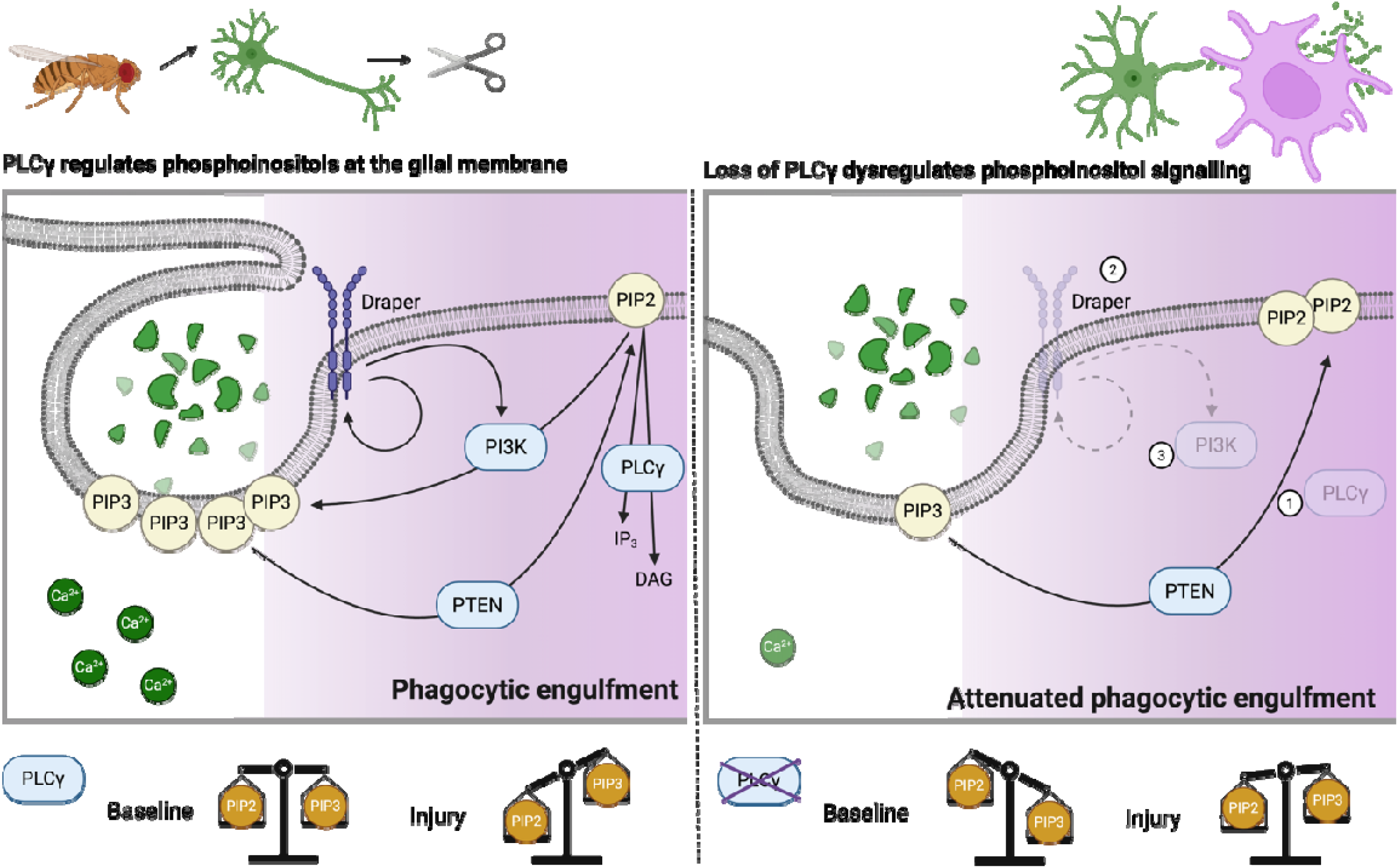

## Introduction

Phosphoinositide signalling is an evolutionarily conserved mechanism that regulates a broad range of cellular processes including membrane trafficking, cytoskeletal remodelling, calcium signalling, receptor organisation and vesicular transport. Tight spatial and temporal regulation of phosphoinositide metabolism is particularly important in the nervous system, where phosphoinositides control neuronal excitability, synaptic transmission and glial responses to injury and disease. Increasing genetic and mechanistic evidence now implicates dysregulated phosphoinositide signalling in neurological disorders spanning neurodevelopmental disease, epilepsy and neurodegeneration (Balla, 2013; Di Paolo & De Camilli, 2006). Several phosphoinositide-metabolising enzymes have been directly linked to neurodegenerative disease pathogenesis. Mutations in the phosphoinositide phosphatase *SYNJ1* cause early-onset Parkinsonism and are associated with Down syndrome-associated neurodegeneration (Choudhry et al., 2021), while mutations in *FIG4*, which regulates PI(3,5)P2 turnover, cause Charcot-Marie-Tooth disease type 4J and have been linked to amyotrophic lateral sclerosis (Lenk et al., 2011; Osmanovic et al., 2017). Beyond monogenic disease, genome-wide association studies have identified phosphoinositide signalling pathways as major contributors to late-onset Alzheimer’s disease (AD) risk. Rare coding variants in *PLCG2* are associated with reduced AD risk, while variants in *INPP5D/SHIP1* increase disease susceptibility (Bellenguez et al., 2022; Kunkle et al., 2019; Lambert et al., 2013).

Among phosphoinositide associated genes implicated in AD, phospholipase C gamma 2 (PLCG2) has emerged as a particularly important microglial signalling molecule. PLCG2 expression is enriched in microglia and is strongly induced in disease-associated microglial states in both human AD tissue and mouse models (Tsai et al., 2023; Zhou et al., 2012). PLCγ enzymes hydrolyse phosphatidylinositol-4,5-bisphosphate (PI(4,5)P2) to generate the second messengers inositol-1,4,5-trisphosphate (IP3) and diacylglycerol (DAG), thereby coupling membrane phosphoinositide turnover to intracellular calcium release and downstream signalling cascades. In peripheral immune cells, PLCG2 functions downstream of immunoreceptors and integrin pathways to regulate phagocytosis, migration and inflammatory signalling (Jackson et al., 2021; Obst et al., 2021). Germline activating mutations in *PLCG2* cause the autoinflammatory syndromes PLAID and APLAID, highlighting the importance of balanced PLCG2 signalling for immune homeostasis (Baysac et al., 2024; Welzel et al., 2022).

In the central nervous system, PLCG2 acts downstream of the microglial receptor TREM2, another major AD risk gene involved in lipid sensing, phagocytosis and injury responses (Obst et al., 2021; Sims et al., 2017). TREM2 activates PLCG2 through DAP12/SYK signalling, promoting intracellular calcium responses, survival and cytoskeletal remodelling in microglia (Hodges et al., 2021; Li et al., 2022; Obst et al., 2021). Functional studies of the AD-protective PLCG2 P522R variant suggest that it acts as a mild hypermorphic allele that enhances microglial immune activity and modifies phosphoinositide dynamics (Maguire et al., 2021; Takalo et al., 2020). Consistent with this, PLCG2 has been implicated in microglial clustering around amyloid plaques, phagocytosis of amyloid material and inflammatory responses in mouse and human models (Andreone et al., 2020; Tsai et al., 2022). However, despite increasing evidence linking PLCG2 to microglial engulfment functions, whether PLCγ signalling directly regulates clearance of degenerating neuronal material following injury remains poorly understood.

Many of the fundamental mechanisms controlling glial engulfment have been elucidated using *Drosophila*, where conserved phagocytic pathways mediate rapid clearance of damaged neurons. The transmembrane receptor Draper, a functional homologue of mammalian MEGF10 and Jedi receptors, is the central engulfment receptor required for glial phagocytosis of severed axons and apoptotic neurons (Freeman et al., 2003; Logan & Freeman, 2007; MacDonald et al., 2006). Following neuronal injury, Draper is activated through the adaptor Ced-6 and downstream Src42A/Shark kinase signalling, leading to Rac1-dependent cytoskeletal remodelling and engulfment (Awasaki et al., 2006; Ziegenfuss et al., 2008, 2012). Orion was found to bridge phosphatidylserine and Draper to facilitate glial recognition of neuronal debris (Ji et al., 2023). Subsequent transcriptional activation through JNK/AP-1 and STAT92E pathways further amplifies Draper expression to sustain phagocytic responses in surrounding ensheathing glia membranes (Freeman & Doherty, 2006; Logan et al., 2012; Lu et al., 2014; Macdonald et al., 2013). PI3K and Stat92E signalling converge to regulate the glial injury response in *Drosophila* (Doherty et al., 2014a) and degradation of engulfed neuronal material through LC3-associated phagocytosis-like mechanisms (Szabó et al., 2023). Although these receptor-proximal signalling pathways have been extensively characterised, considerably less is known about how membrane lipid signalling and phosphoinositide turnover contribute to Draper-mediated engulfment.

Phosphoinositide metabolism is highly conserved between flies and mammals, providing an opportunity to dissect how phosphoinositide signalling regulates glial function in vivo. Here, we investigate the role of *small wing* (*sl*), the *Drosophila* orthologue of human PLCγ genes including *PLCG2*, in glial engulfment and AD-relevant pathology. Using injury paradigms and genetically encoded phosphoinositide reporters, we show that neuronal injury induces dynamic remodelling of glial PIP2 and PIP3 downstream of Draper activation. Loss of *sl* disrupts these phosphoinositide responses, impairs Draper upregulation and delays engulfment of degenerating axons. In addition, manipulation of *sl* and expression of human *PLCG2* variants modifies organismal survival in a *Drosophila* Aβ42 model. Together, our findings identify PLCγ-dependent phosphoinositide turnover as a conserved regulator of Draper-mediated glial engulfment and provide mechanistic insight into how altered PLCG2 signalling may contribute to neurodegenerative disease-associated glial states.

## Methods

### *Drosophila* maintenance and Stocks

Experimental crosses were maintained in temperature-controlled incubators at 18°C, 25°C or 29°C under a 12 h light/dark cycle. The following fly stocks were obtained from the Bloomington Drosophila Stock Center (BDSC; NIH P40OD018537): *repo-Gal4* (RRID:BDSC_7415), *Tubulin-Gal4* (RRID:BDSC_5138) (Lee & Luo, 1999), *5xUAS-aos::Aβ42Arc* (RRID:BDSC_33773), *5xUAS-LacZ* (RRID:BDSC_8530), *slCRIMIC* (RRID:BDSC_81213), *5xUAS-mCD8::GFP* (RRID:BDSC_27398), *5xUAS-PLCδ-PH::GFP* (RRID:BDSC_39693) (Várnai & Balla, 1998), *GRP1-PH::GFP* (RRID:BDSC_8163) (Britton et al., 2002), *UAS-GFP-Valium10* (RRID:BDSC_35786), *CaryPattP2* (RRID:BDSC_36303), and *20xUAS-GCaMP6f* (RRID:BDSC_42747). The following RNAi lines were obtained from the Vienna Drosophila Resource Center (VDRC): *5xUAS-slRNAi* (RRID:BDSC_108593, RNAi I; RRID:BDSC_32906, RNAi II), *5xUAS-DraperRNAi* (RRID:BDSC_27086), and *5xUAS-ptenRNAi*. Olfactory receptor neurons (ORNs) were labelled using *OR85e-mCD8::GFP* (Couto et al., 2005).

### Longevity

24 hour mated females were selected from the F1 progeny and separated at a density of 10-15 flies per vial, containing feeding media. At least 10 vials per genotype were collected (total of ∼100 flies per genotype) and maintained at 29°C. Flies were flipped into fresh feeding media every 2-3 days and number of deaths were recorded at each passage. Survival data was presented as Mantel-Cox survival curves and log rank tests were performed for statistical comparisons between genotypes.

### Rapid Iterative Negative Geotaxis (RING)

All RING assays were performed in the same environment at 22-25 °C and flies were left to acclimatise within the apparatus for 10 mins before starting the assay. Up to 20 flies were transferred into a single clear polystyrene 2cm diameter vial and sealed with a plug that sat flush with the top of the vial. Up to six vials were loaded into a custom-built holding frame, which moves vertically along a metal rail. To initiate negative geotaxis, the frame was released once from a defined height of 20cm, to tap down all flies. As the flies ascended the vial, their position was photographed every second for 10 seconds. A single RING assay trial consisted of one round of initiating negative geotaxis, followed by digitally capturing their locomotive behaviour over 10 seconds. The resulting images were used to determine the vertical position of the flies within the vial. For all experiments, the distance travelled was averaged over 5 consecutive RING trials, separated by 30 seconds of rest. Digital images (.JPG) of the flies were processed using the FIJI software (ImageJ). The full FiJi script is available via Github (https://github.com/Fly-Cardiff/Fly-RING-Analysis)

### Molecular Cloning and Fly Stock generation

To generate site directed insertion fly lines, cDNA encoding the *Drosophila sl* gene (DGRC, GOLD RE62235) and human *PLCG2* gene (DGRC, HSCD00506018) were subcloned into the *5xUAS-pJFRC5* vector (Gifted from Gerald Ruben, Addgene, plasmid 26218). *sl* and *PLCG2* cDNA were amplified by PCR using primer sequences: *sl* forward TCAGCGGCCGCACAACCAAAATGAGCTGCTTTAGTGCGAT, *sl* reverse GTATCTAGACTACGGTGCGGTAACATTTG, *PLCG2* forward TCAGCGGCCGCACAACCAAAATGTCCACCACGGTCAATGT, *PLCG2* reverse GTATCTAGACTACGGTGCGGTAACATTTG that added NotI and XBaI restriction sites to the 5’ and 3’ ends. Amplified PCR product was isolated from agarose gels according to manufacturer’s instructions (Thermo Fisher, K220001). 1µg of *5xUAS-pJFRC5* vector, *sl* and *PLCG2* DNA were digested with *Not1-HF* and *XBaI* restriction enzymes and ligated using T4 DNA ligase (New England Biolabs, M0202S), with addition of alkaline phosphatase to prevent re-ligation of the vector backbone. DH5a cells (Thermo Fisher, 18263021) were transformed with ligation product and individual colonies were picked from LB + Ampicillin resistance plates. Plasmid DNA from positive colonies was isolated using the QIAprep Spin Miniprep kit (Qiagen, 27106) and sequence verified (Genewiz). For site-directed mutagenesis to generate *PLCG2-R552*, 4µg of plasmid containing wildtype *PLCG2-P552* DNA was provided to commercial vendor Genscript (Netherlands, Leiden). Site-directed mutagenesis introduced a C to G base substitution at position 1564bp of the *PLCG2* insert, resulting in a Proline to Arginine conversion.

### *Drosophila* embryo injection for transgenic production

Embryo injection and site directed integration of verified *sl* and *PLCG2* variant plasmids was conducted by a commercial vendor (BestGene (USA). Variant plasmids were injected in to *Drosophila melanogaster* embryos harbouring attP40 (BDSC 36304) and attP2 (BDSC 8622) landing sites, with constructs integrated using PhiC31 integrase-mediated site-specific recombination. An empty JFRC vector control line was generated in parallel and used as a transgenic insertion control throughout the study. Successful transformants were identified, back-crossed into a *w1118* background, and balanced prior to experimental use.

*RNA Sequencing Small wing* expression was reduced in glial cells using the *repo-GAL4* driver and a sl targetting RNAi (BDSC 3290). At 14 days, RNA from 10 heads per sample, was extracted using the RNAqueous-Micro kit (Thermo Fisher Scientific, AM1931). Thereafter, TruSeq mRNA libraries were generated (Illumina, 20020595) by the Cardiff University Genomics Research Hub and sequenced, at a depth of 6.5 million reads, with an Illumina NextSeq500. Quality was assessed using FastQC (0.11.8) subsequently, adapters, tails, individual low-quality bases (phred <3), average low-quality four base regions (phred <15) and truncated reads (<20 bases, Trimmomatic (Bolger et al., 2014), RStudio 2022.12) were removed. Cleaned reads were then aligned to the *Drosophila* BDGP6.32 genome using STAR (genomeGenerate, alignReads (Dobin et al., 2013). RStudio 2022.12) and counted featureCounts (Liao et al., 2014), RStudio 2022.12. Non-coding RNA, transposons and unassigned reads were removed using SARTools (Varet et al., 2016) to concentrate the analysis solely on protein coding mRNA. After variance stabilization, principal component analyses (prcomp, R), heat maps and dendrograms were computed (pheatmap, RStudio 2022.12). Upon evaluation of these dimensionality reductions, one control replicate (C4) was excluded as an outlier. Finally, differential gene expression was performed using DESeq2 (Love et al., 2014). Differentially expressed genes were defined as those with an adjusted p-value less than 0.05 and a log2 foldchange greater than +/−0.58. All genes showing differential expression, regardless of the direction of change, were entered into g:Profiler for analysis to identify significantly enriched pathways and gene ontology terms, graphed on SRPlot. Data is accessible on ArrayExpress (E-MTAB-14080) and the analysis pipeline is deposited on GitHub (https://github.com/Fly-Cardiff/RNA_Seq).

### Olfactory Receptor Neuron Injury

Neuronal injury was induced by bilateral ablation of maxillary palps or third antennal segments of adult flies (7-14 days old). This was achieved using fine forceps (Dumont, carbon forceps) under a dissection microscope, following standard procedure as described in previous studies (MacDonald et al., 2006; Vosshall et al., 2000).

### Fluorescent immunohistochemistry

Adult heads (7-14 days old) were removed and fixed in 4% paraformaldehyde in PBS (4% PFA) for 16 min at room temperature. After washing them with (0.1% Triton X-100 in PBS (PBST) five times, brains were dissected in PBST using forceps and fixed a second time with 4% PFA for 16 min at room temperature. Fixed brains were blocked in 10% goat serum/PBST for 1 hr at room temperature and incubated in primary antibodies in blocking solution at 4°C overnight. The following day, brains were washed with PBST and incubated in secondary antibodies for 2 hrs at room temperature. Brains were washed five times with PBST, mounted on glass slides in VECTASHIELD Mounting Medium (Vector Labs) and stored at 4°C for imaging.

### Antibodies

Primary antibodies used in this were Rabbit anti-GFP (Thermo Fischer A11122, 1:1000 for IHC), Mouse anti-Draper (Developmental Studies Hybridoma Bank (DSHB) 8A1, 1:50 for IHC); anti-Aβ (1-16, 6E10, mouse, 1:400, Biolegend #803001); Mouse anti-repo (DSHB 8D12, 1:100 for IHC). Secondary antibodies were Alexa FluorTM 488, Donkey anti-rabbit (Thermo Fischer Scientific A21206, 1:500 for IHC), Alexa FluorTM 568 and Goat anti-Mouse (Thermo Fischer Scientific A11004, 1:500 for IHC).

### Confocal microscopy and image analysis

A Zeiss Cell Observer spinning disk confocal microscope was used for all brain imaging, with onfocal microscopy settings kept constant throughout each set of experiments. For fluorescence quantification, whole z-stack projections were used and the mean grey value of specific structures measured in ImageJ (National Institute of Health). Following neuronal injury, pixel intensity was measured by tracing glomeruli and axon regions separately and normalising to background levels of GFP. For Draper immunofluorescent staining analysis, measurements were taking from the antennal lobe border where signal was most concentrated. For genetically encoded fluorescent reporters GCaMP6f, PLCδ-PH::GFP and GRP1-PH::GFP, adult brains were dissected and fixed as described, washed with PBST five times and mounted in VECTASHIELD for imaging. Measures of fluorescence encompassed the entire optic lobe.

### Electrochemiluminescence assay for Aβ_42_

40 fly heads per genotype (20 male and 20 female) at 7 days were homogenised in 150µl of soluble extraction buffer (50 mM HEPES pH 7.3, 5 µM EDTA and protease inhibitor) (Complete mini EDTA-free protease inhibitor), vortexed briefly and incubated for 10 mins at room temperature. Tissue homogenate was sonicated in a 4°C water bath for 4 mins following a 30 second on/off cycle. The homogenate was then centrifuged at 17,000 x g for 5 mins at 4°C and supernatant transferred to a fresh 1.5 ml microcentrifuge tube, for the soluble fraction of Aβ_1-42_. The remaining pellet was re-homogenised with 55µl of insoluble extraction buffer (50 mM HEPES pH 7.3, 5 µM EDTA, 5 mM Guanidinium Hydrochloric acid and protease inhibitor), vortexed briefly and incubated for 10 mins at room temperature. The homogenate was sonicated in a water bath set to 4°C for 4 mins following a 30 second on/off cycle and then centrifuged at 17,000 x g for 5 mins at 4°C. The collected supernatant comprised of the insoluble fraction.

Soluble and insoluble Aβ peptides (38,40,42) were measured using Mesoscale Discovery V-PLEX Aβ peptide panel 1 (6E10) kit (K15200E) or the V-PLEX Aβ42 peptide (4G8) kit (K150SLE) according to manufacturer’s instructions. Soluble and insoluble Aβ extractions were diluted 1 in 5 and 1 in 10 respectively in Diluent 35. Wells of the MSD plates were blocked using 150µl Diluent 35 at room temperature for 1 hour, with gentle rocking. After blocking, wells were washed and incubated for 2 hours with 25µl of detection antibody solution and 25µl of prepared samples, Aβ calibrators or controls per well, plated in duplicate. After sample incubation, the plate was washed 3 times with MSD wash buffer and 150µl of 2x Read Buffer T was added to each well, before taking a recording of analyte levels on the MESO QUICKPLEX SQ120. Analysis of Aβ peptide was performed using the MSD Workbench 4.0 software. Aβ peptide concentration was normalised to the total protein concentration for each sample as determined by BCA assay.

### Statistical analysis

Unless stated otherwise, GraphPad Prism (version 9.0) was used for statistical analysis. 2-tailed t-tests were used for comparisons between two groups, one-way ANOVA or two-way ANOVA for comparing multiple groups, with multi-comparison tests as indicated in the figure legends. Significant differences (p-value less than 0.05) are marked with asterisks. All graphs show the mean with standard error of the mean (SEM). The exact number of samples used in each experiment is provided in the legends of corresponding figures.

## Results

### Small wing contributes to normal function of *Drosophila* glia

In human microglia, PLCG2 plays important roles in activating downstream signalling processes in recognition of external ligands by receptors including TREM2 and TLRs. sl is the fly ortholog of PLCG1 (39% identify, 57% similarity) and PLCG2 (37% identify, 57% similarity). Global RNAi-mediated knockdown of *sl* caused the hallmark phenotype of small wings recapitulating the mutant phenotype and caused a 50% reduction of mRNA compared to control (p<0.001), (Supplementary material, Fig. 1). In the central brain sl is expressed in glia (Fig. 1A) and knocking down *sl* specifically in glia causes significant transcriptional changes (Fig. 1B).

**Figure 1.**
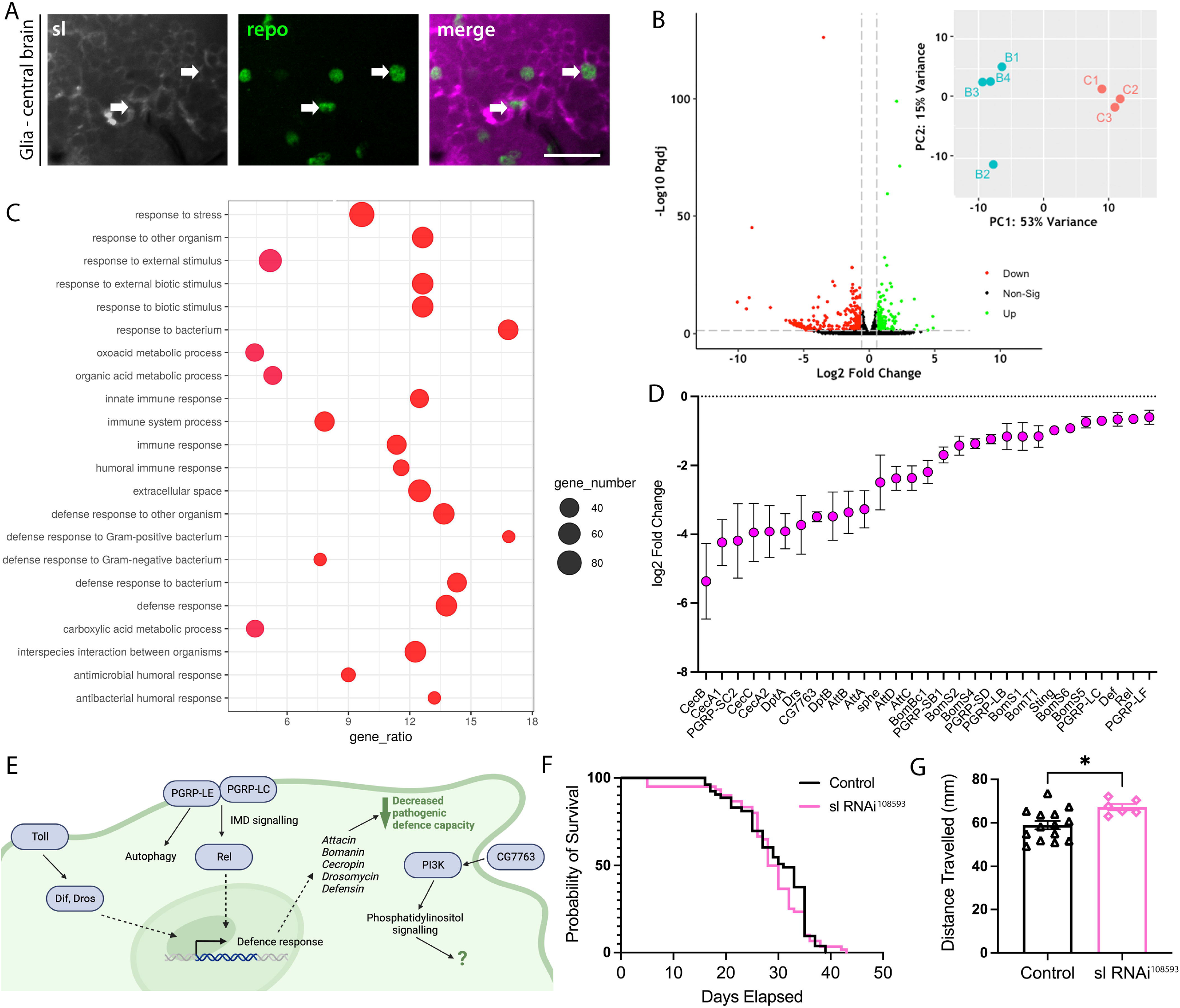
Small wing regulates glial signalling in *Drosophila*. A) Endogenous expression of *sl* is ubiquitously expressed in the brain and present in glial cells. *sl* CRIMIC promoter-trap driving expression of membrane-tethered mCD8::RFP (magenta) and immunofluorescent staining for glia specific nuclear repo (green). B) RNAi-mediated knockdown of sl in glia causes differential gene expression changes in the brain and a clear separation of genotypes by PCA. C) Gene Ontology analysis of differentially expressed genes indicates significant enrichment for several biological pathways associated with immunity and cellular defence against pathogens. D) Most significantly downregulated genes from immunity-related pathway. E) Graphical model of intracellular signalling changes identified and putative mechanism following glial specific *sl* knockdown. F) Knockdown of *sl* in glia had no effect on longevity compared to control (n=53-90 per group) however, G) presented a modest improvement of locomotor behaviour compared to control (n=6-14 vials per group). Data was analysed using Log Rank Mantel-Cox test or One-way ANOVA with Tukey’s multiple comparison test and statistical differences are shown as: ****p<0.0001 and *p<0.05. Error bars represent ±SEM. Scale bar = 10µM.

Control and experimental samples clustered distinctly with principal components 1 and 2 accounting for 54% and 15% of variance, respectively. Differential gene expression analysis revealed 163 significantly upregulated and 258 downregulated genes after *sl* knockdown compared to control (Fig. 1B). Gene ontology enrichment analysis of these differentially expressed genes (irrespective of directional change) revealed a significant enrichment of immune pathways including innate immunity, antibacterial response and defence responses to gram-positive and -negative bacteria (Fig. 1C). Notable downregulated mechanisms indicated misregulation of innate immune IMD pathway components, including peptidoglycan recognition receptors (PGRP-LC, -LE), NFkB (Rel) and antimicrobial peptides (attacin, bomamin, cecropin, drosomycin). AMPs downstream of Toll receptor signalling were also found to be down regulated (Defenin, Drosomycin) (Fig 1D-E). CLEC3A (CG7763) was also significantly downregulated (Fig. 1D-E), which has been shown to interact with phosphatidylinositol 3-kinase (PI3K) in several cancer models (Chen et al., 2022; Ni et al., 2018), where it is required for PIP3 synthesis from PIP2. PI3K is involved cellular functions such as proliferation, cell growth, differentiation, motility, survival and intracellular trafficking and therefore sl-CG7763-PI3K signalling could also impact specialized functions of glial cells. Notably, in *Drosophila* glia, PI3K and Stat92E signalling pathways regulate responsiveness to axonal injury by modulating expression of the engulfment receptor Draper (Doherty et al., 2014a).

Despite inducing immune-related glial responses, *sl* knockdown had no significant detrimental effect on survival (Fig. 1F). Locomotor performance was modestly increased relative to controls (p<0.05, Fig. 1G), consistent with previous reports demonstrating that constitutive *sl* null mutants are viable and display minimal overt phenotypes. (Schlesinger et al., 2004; Thackeray et al., 1998).

### Glia lacking sl are deficient in normal engulfment activity

A key response to bacterial infection and neuronal injury is the ability of glia to recognise and engulf pathogens and damaged tissues, a process in which mammalian PLCG2 contributes (Maguire et al., 2021; Takalo et al., 2020). Following bilateral ablation of the sensory antennae, axons of olfactory receptor neurons projecting into the *Drosophila* central brain are severed inducing Wallerian degeneration (Ziegenfuss et al., 2008). Fragments of degenerating olfactory neuron (ORN) axons are effectively cleared by ensheathing glia, which surround olfactory glomeruli in the adult brain (Logan et al., 2012).

To test the contribution of *sl* in this process, we performed adult glia RNAi knock down in flies expressing a GFP reporter restricted to the Odorant receptor 85e (OR85a) subset of ORN neurons. These neurons have a stereotyped morphology in the antennae and olfactory glomeruli, providing a tractable system for measurement of glial clearance of severed axon and synapse debris. Following bilateral ablation of antennae, ORN85a axons positive for GFP in control flies were efficiently cleared from the central brain within 3 days, with synaptic debris within the glomeruli cleared by 5 days (Fig 2A). In flies with reduced glial *sl*, GFP positive OR85a axonal fragments were observed within olfactory glomeruli for up to 14 days post injury, with small fragments of axon also still present up to 3-days post-injury (Fig. 2A, Supplementary material Fig. 2). At 3 days, 50% of axon debris failed to be engulfed compared to control (p<0.01) indicating a delay of engulfment (Fig. 2B). Equally glomeruli engulfment was strongly inhibited even at 14 days post injury (p<0.0001), (Fig. 2C).

**Figure 2.**
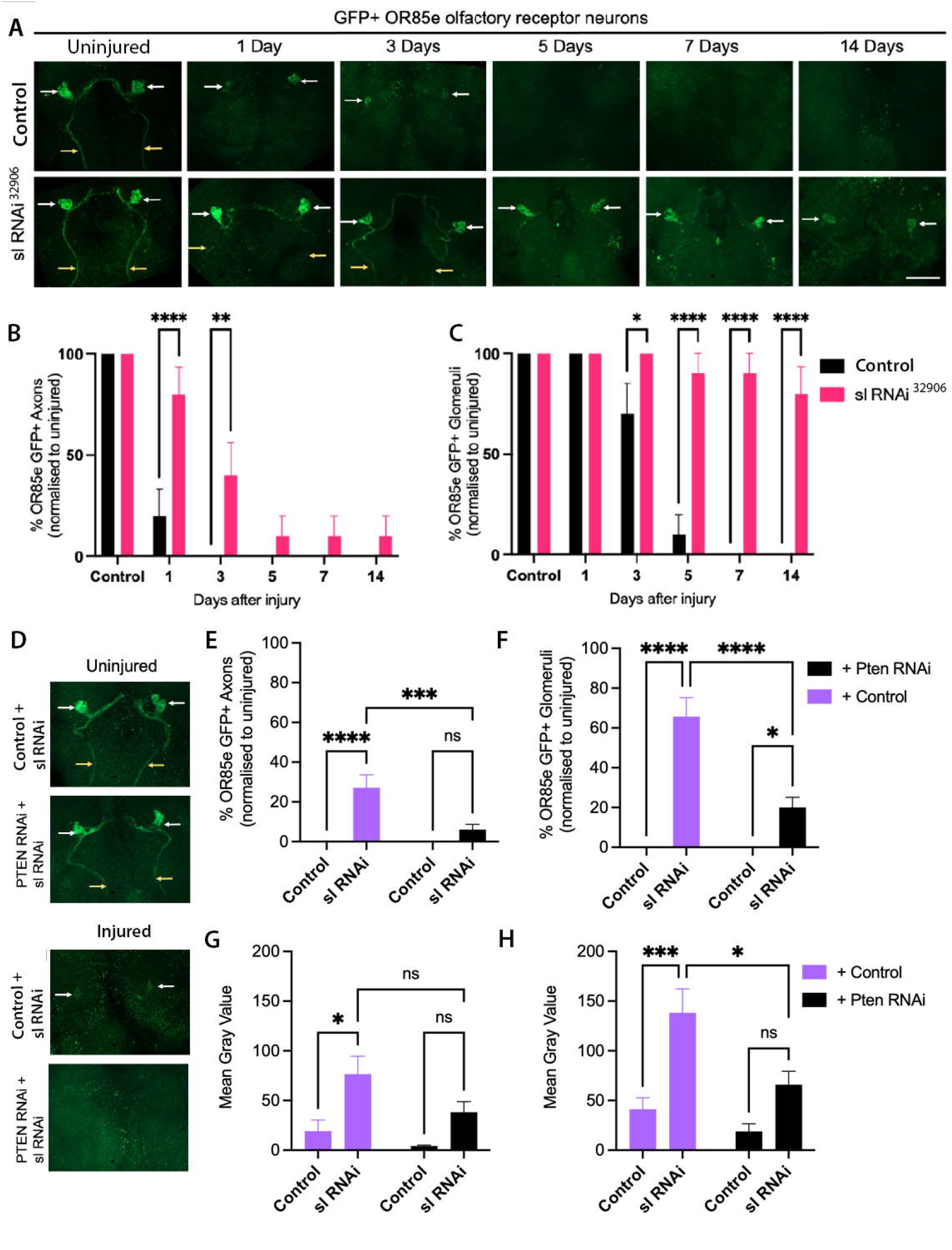
Small wing contributes to glial engulfment of neuronal debris. A) Timecourse of GFP-labelled maxillary olfactory receptor neurons following axotomy in control (attP2) or glial driven *sl* RNAi expressing flies. B) Analysis of percentage of antennal lobes containing visible GFP+ axons derived from severed OR85e maxillary ORN neurons and C) GFP+ glomeruli detected significantly delayed clearance in *sl* RNAi expressing flies. D) Representative images of GFP-labelled maxillary ORN neurons before and 5 days after injury, showing axons (yellow arrows) and glomeruli (white arrows), showing suppression of engulfment deficit by combined glial specific RNA-mediated knock down of *sl* and *Pten*. E) Analysis of percentage of antennal lobes containing GFP+ axons and F) GFP+ glomeruli across experimental lines. G) Analysis of mean GFP fluorescence of OR85e axons and H) glomeruli across for combined *sl* and *Pten* knock down. Data was analysed using Kruskal Wallis test with Dunn’s multiple comparison test or a one-way ANOVA and Dunnett’s multiple comparison test. Graphs show mean±SEM, *p<0.05, **p<0.01, ***p<0.001 and ****p<0.0001, n=7-10 brains per group. Scale bars = 50µm.

To investigate whether phosphoinositide signalling regulates glial engulfment downstream of PLCγ activity, we examined the genetic interaction between *sl* and the lipid phosphatase *pten* following axonal injury. PTEN is a lipid phosphatase that antagonises PI3K signalling by converting phosphatidylinositol (3,4,5)-trisphosphate (PIP3) back to phosphatidylinositol (4,5)-bisphosphate (PIP2), thereby limiting downstream signalling pathways that regulate membrane dynamics, cytoskeletal remodelling and phagocytic activity. Simultaneous knock down of *sl* and *pten* specifically in glial cells rescued the delayed engulfment phenotype observed in *sl* knockdown animals (Fig. 2D), restoring more rapid clearance of degenerating neuronal material by glia. While pten knockdown in glia did not change the engulfment profile of wild-type neurons, combined glial *pten* and *sl* knockdown significantly decreased the number of brains with GFP debris remaining in the axons (p<0.001) and the glomeruli (p<0.0001) regions and the GFP fluorescence intensity of the glomeruli region (p<0.05), (Fig 2D-H) indicating a rescue effect. These findings suggest that loss of *pten* elevates PIP3 levels, thereby compensating for reduced PLCγ activity and promoting engulfment. Together, these data support a model in which phosphoinositide signalling is a key regulator of glial phagocytic responses following neuronal injury.

### Glial phosphoinositide dynamics in response to neuronal injury

To further investigate the role of phosphoinositides in glial engulfment of neuron debris, we next used genetically encoded fluorescent probes to assess changes their glial distribution following axonal injury. To examine the glial distribution of PIP2, we expressed a GFP-tagged pleckstrin homology (PH) domain of PLCd (*PLCd-PH::GFP*) (Várnai & Balla, 1998), specifically in glial cells. Following bilateral antennal ablation, glial PIP2 reporter fluorescence progressively declined over the first 48 h after injury (Fig. 3A,B), reaching its lowest levels at 48 h post-ablation (p<0.001). By 5 days post-injury, PIP2 reporter intensity had recovered to levels slightly above baseline (p<0.01; Fig. 3A,B).

**Figure 3:**
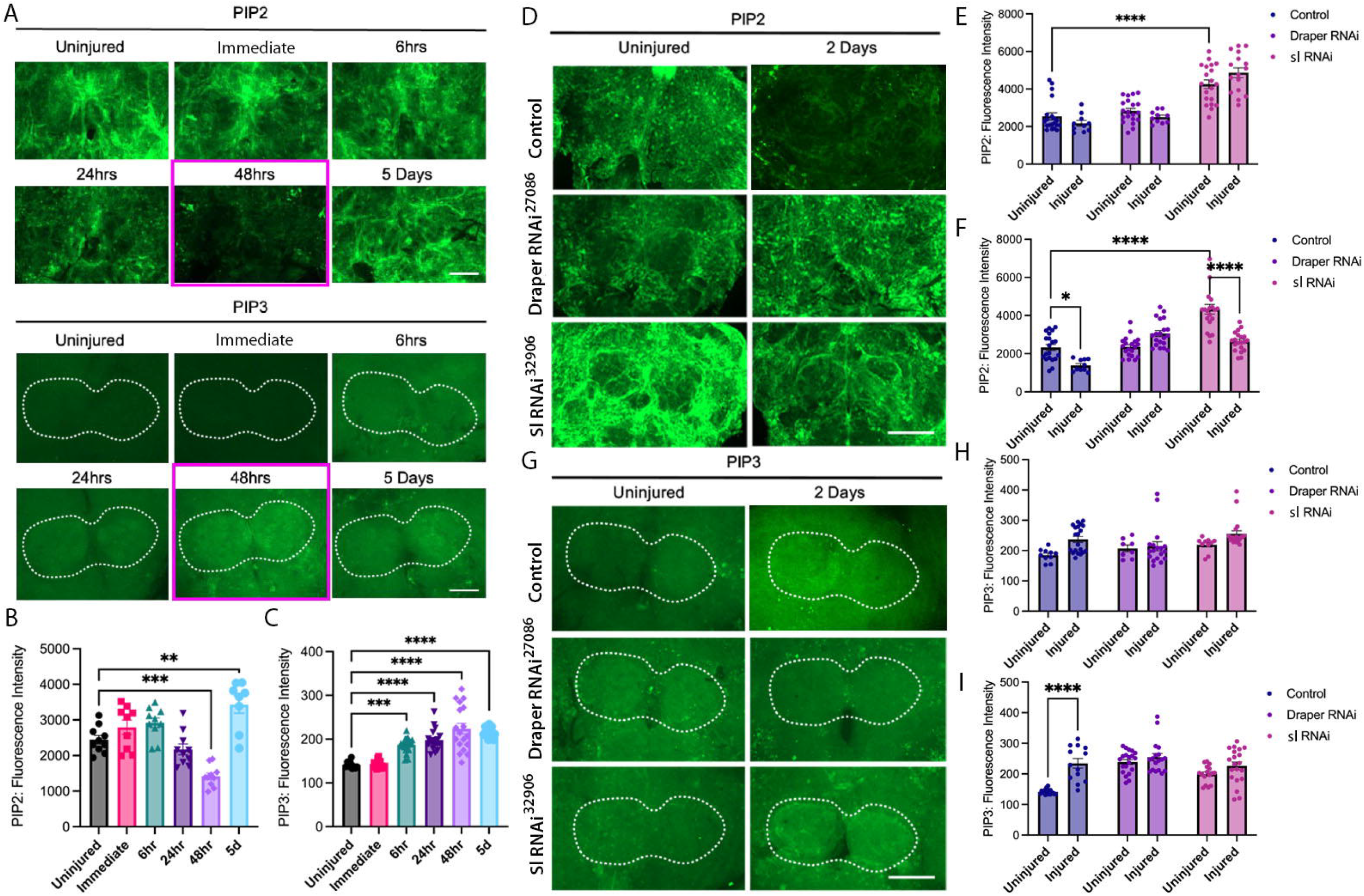
Glial phosphoinositide dynamics in response to neuronal injury is altered by loss of small wing. A) Representative images of Repo-Gal4 driven PIP2 (*PLCδ-PH::GFP*) and ubiquitous expressed PIP3 (*GRP1-PH::GFP*) fluorescent reporter brains before, immediately post-injury, 6hrs, 24hrs, 48hrs and 5 days after injury in LacZ-expressing flies. B) Quantification of PIP2 and C) PIP3 fluorescent reporters in antennal lobe before and after antennal nerve injury. D) Representative images of Repo-Gal4 driven PIP2 reporter brains before and 48hrs after injury in sl and Draper RNAi expressing flies. E) Quantification of PIP2 fluorescent reporter before and immediately following (1-5 mins) antennal lobe injury and F) after 48hrs. G) Representative confocal z-stacks of PIP3 reporter brains before and 48hrs after injury in Draper and sl RNAi expressing flies. H) Quantification of PIP3 fluorescent reporter in antennal lobe before and immediately following (1-5 mins) antennal nerve injury and I) after 48hrs. Statistics was performed with either One-way ANOVA and Dunnett’s multiple comparison test or Two-way ANOVA with Tukey’s multiple comparison test. Graphs show mean±SEM, **p<0.01, ***p<0.001, ****p<0.0001, n=8-10 brains per group. Scale bar = 50µm.

To assess PIP3 dynamics, we utilised a ubiquitously expressed GFP reporter fused to the PH domain of Cytohesin 3 (*GRP1-PH::GFP*) (Britton et al., 2002). In contrast to PIP2, PIP3 reporter fluorescence increased within the olfactory glomeruli region containing ensheathing glia actively engaged in debris clearance from as early as 6 h post-injury (Fig. 3A,C). PIP3 signal continued to rise throughout the injury response, reaching maximal levels between 2 and 5 days post-ablation (p<0.0001; Fig. 3A,C).

As glial levels of phosphoinositides were found to actively change in response to neuronal injury, we next assessed if slowing engulfment altered their regulation. In flies lacking Draper, phosphoinositide levels in the olfactory glomeruli region were quantified at uninjured baseline and two days after bilateral antennal ablation. In contrast to controls, PIP2 (Fig 3D-F) and PIP3 (Fig 3G-I) intensity remained stable in flies with reduced glial *Draper* expression before and after injury, indicating membrane phosphoinositide integration changes occurs downstream of ligand recognition and initiation of phagocytic engulfment.

We next investigated whether *sl* contributes to the regulation of glial membrane phosphoinositides and their dynamic redistribution during engulfment activation. Prior to injury, glial PIP2 levels were significantly elevated following *sl* knockdown compared with controls (p<0.0001; Fig. 3D–F), whereas basal PIP3 levels remained unchanged (Fig. 3G–I). Immediately following antennal ablation (<1 min), membrane-associated PIP2 and PIP3 levels were largely unchanged across all genotypes, indicating that acute injury alone does not rapidly alter phosphoinositide distribution (Fig. 3E,H).

At 48 h post-injury, control glia displayed the expected reduction in membrane-associated PIP2, consistent with active phosphoinositide turnover during engulfment (p<0.0001; Fig. 3F). In contrast, Draper knockdown prevented this injury-induced depletion of PIP2, while *sl*-deficient glia exhibited only a partial reduction, with PIP2 levels remaining significantly elevated relative to controls (Fig. 3F). These findings suggest that effective engulfment requires substantial depletion of membrane-associated PIP2 and that *sl* activity contributes to this process.

In parallel, injury-induced accumulation of membrane-associated PIP3 was markedly impaired in both Draper- and *sl*-depleted glia at 48 h post-injury (Fig. 3I), despite normal basal levels prior to injury. Together, these findings indicate that Draper and *sl* function within a shared signalling pathway that coordinates phosphoinositide remodelling during glial engulfment, linking PLCγ activity to the membrane lipid transitions required for efficient clearance of neuronal debris.

### Small wing promotes upregulation of Draper and Ca^2+^ in response to axon severing Injury

In response to axonal injury, glia upregulate the engulfment receptor Draper to initiate clearance of neuronal debris. Draper signalling is further amplified through a positive feedback mechanism that promotes efficient and complete engulfment, with disruption of this pathway resulting in delayed or suppressed debris clearance (Doherty et al., 2014b; Logan et al., 2012; Lu et al., 2017). We therefore investigated whether the impaired glial engulfment observed following *sl* depletion was associated with altered Draper signalling in response to neuronal injury. Consistent with previous reports, antennal axotomy induced a significant increase in Draper levels within ensheathing glia surrounding the olfactory glomeruli, accompanied by extension of glial processes into the injured region (p<0.01; Fig. 4A,B). RNAi-mediated knockdown of Draper abolished this injury-induced signal, confirming the specificity of Draper immunolabelling (Fig. 4A,B). Glial knockdown of *sl* did not alter basal Draper abundance in uninjured flies; however, unlike controls, *sl*-deficient glia exhibited only minimal Draper upregulation following antennal ablation. These findings suggest that *sl* is required for efficient injury-induced amplification of Draper signalling and that impaired Draper upregulation contributes to the reduced engulfment capacity observed following *sl* depletion.

**Figure 4:**
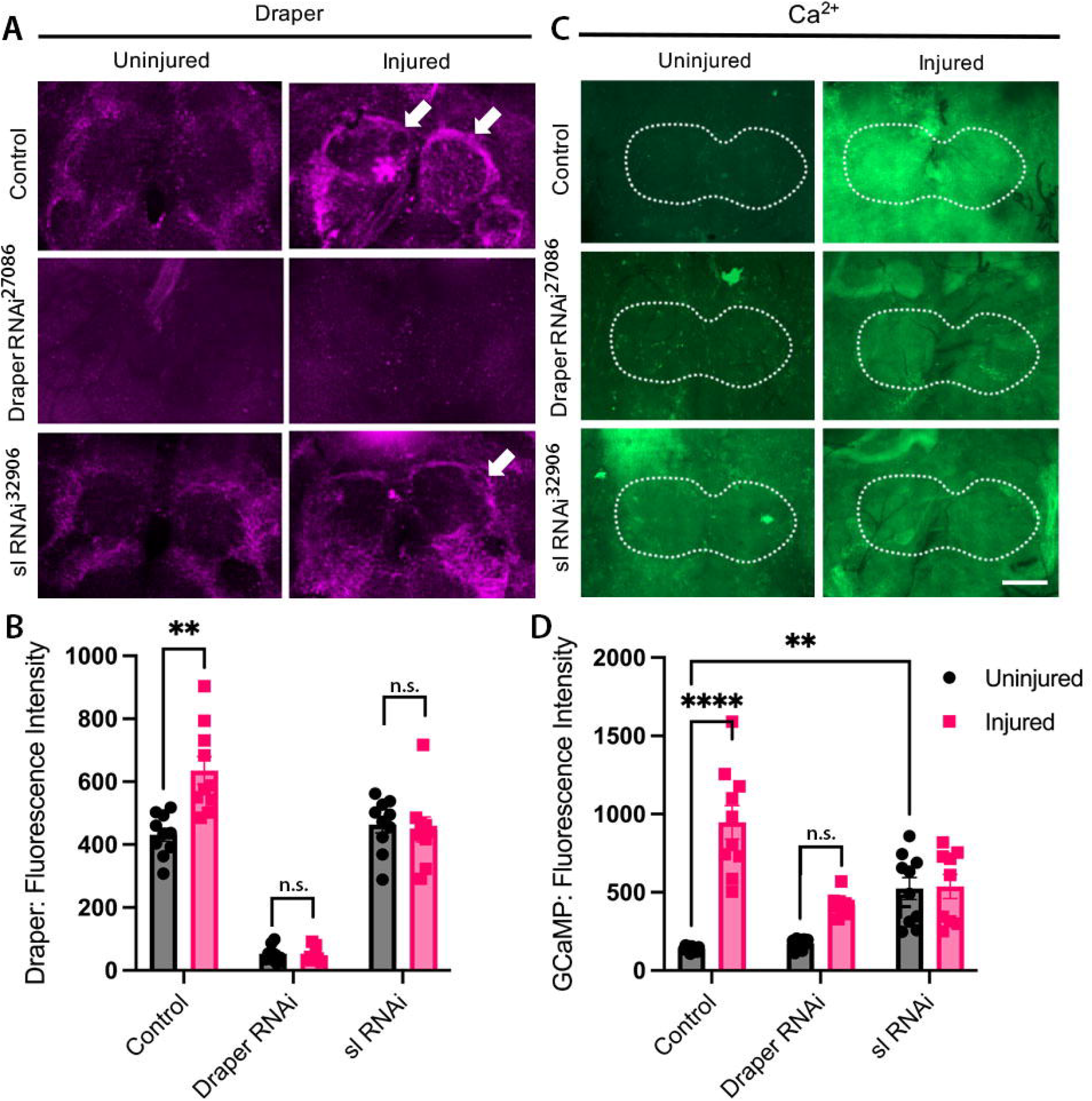
Small wing promotes upregulation of Draper and Ca^2+^ in response to axon severing Injury. A)Immunostaining demonstrating increased Draper levels in antennal lobe in response to antennal severing injury in w-control (white arrows), which is attenuated in flies expressing sl RNAi. Flies expressing Draper RNAi show absence of immunostaining 1 day after injury. B) Quantification of Draper immunostaining fluorescent intensity in antennal lobe of uninjured and injured flies. C) Fluorescent GCaMP6f reporter demonstrates significantly increased Ca^2+^ levels immediately following antennal ablation injury, which was reduced following sl knockdown. D) Flies expressing RNAi targeting Draper or sl had an attenuated GCaMP6f reporter response following injury. Statistics was performed using a Two-way ANOVA with Tukey’s multiple comparison tests. Graphs show mean±SEM, *p<0.05, **p<0.01, ***p<0.001, ****p<0.0001, n=7-10 brains per group. Scale bar = 50µm.

Previous studies have shown that Draper signalling activates Src and Shark-dependent pathways to coordinate glial responses to neuronal injury, including cytoskeletal remodelling and engulfment progression (Lu et al., 2017). In addition, Draper-dependent inflammatory priming in *Drosophila* macrophages has been shown to require calcium-induced JNK signalling, linking Ca2+ mobilisation to enhanced phagocytic responsiveness following corpse engulfment (Weavers et al., 2016). Given that PLCγ enzymes couple receptor activation to intracellular Ca2+ mobilisation through hydrolysis of PIP2, we next investigated whether *sl* regulates injury-induced calcium signalling downstream of Draper during glial engulfment.

We expressed a cytosolic calcium-sensitive GCaMP6f reporter in glia and quantified fluorescence following antennal ablation. Immediately after injury, Ca2+ levels in glia surrounding the olfactory glomeruli increased markedly (p<0.0001; Fig. 4C,D). In contrast, glial Draper knockdown flies exhibited normal basal Ca2+ levels but failed to mount a significant injury-induced calcium response, consistent with impaired engulfment signalling.

Similarly, injury-induced Ca2+ elevation was abolished in *sl*-deficient glia, indicating disruption of downstream calcium mobilisation. Together, these findings demonstrate that *sl* promotes Draper-mediated glial engulfment through regulation of injury-induced calcium signalling.

### Small wing maintains normal phosphoinositide levels in an adult fly model of amyloid pathology

Rare coding variants in *PLCG2*, the human orthologue of *sl* that is highly enriched in microglia, are significantly associated with late-onset Alzheimer’s disease (Bellenguez et al., 2022). In particular, the *PLCG2* P522R coding variant, predicted to function as a hypermorphic allele (Magno et al., 2019), is associated with reduced risk of late-onset AD (Sims et al., 2017). We therefore investigated whether altered glial phosphoinositide regulation influences amyloid-associated pathology in vivo. To model amyloid accumulation in the adult brain, we used *repo-Gal4* to drive expression of the Arctic variant of Aβ_42_ fused to an Argos secretion signal peptide, promoting extracellular amyloid deposition within the central nervous system.

We first examined whether Aβ_42_ expression altered phosphoinositide homeostasis in flies with reduced glial *sl* expression. Glial knockdown of *sl* significantly reduced PIP2 levels in Aβ_42_-expressing brains relative to Aβ_42_ controls (p<0.05; Fig. 5A,B), whereas PIP3 levels were significantly elevated throughout the adult brain (p<0.05; Fig. 5C,D). These findings suggest that *sl* depletion disrupts phosphoinositide balance in the presence of amyloid pathology, consistent with altered phosphoinositide turnover and downstream lipid signalling. Despite these changes, glial *sl* knockdown had no significant effect on levels of either soluble or insoluble Aβ_42_ measured by electrochemiluminescent immunoassay (Fig. 5E,F).

**Figure 5:**
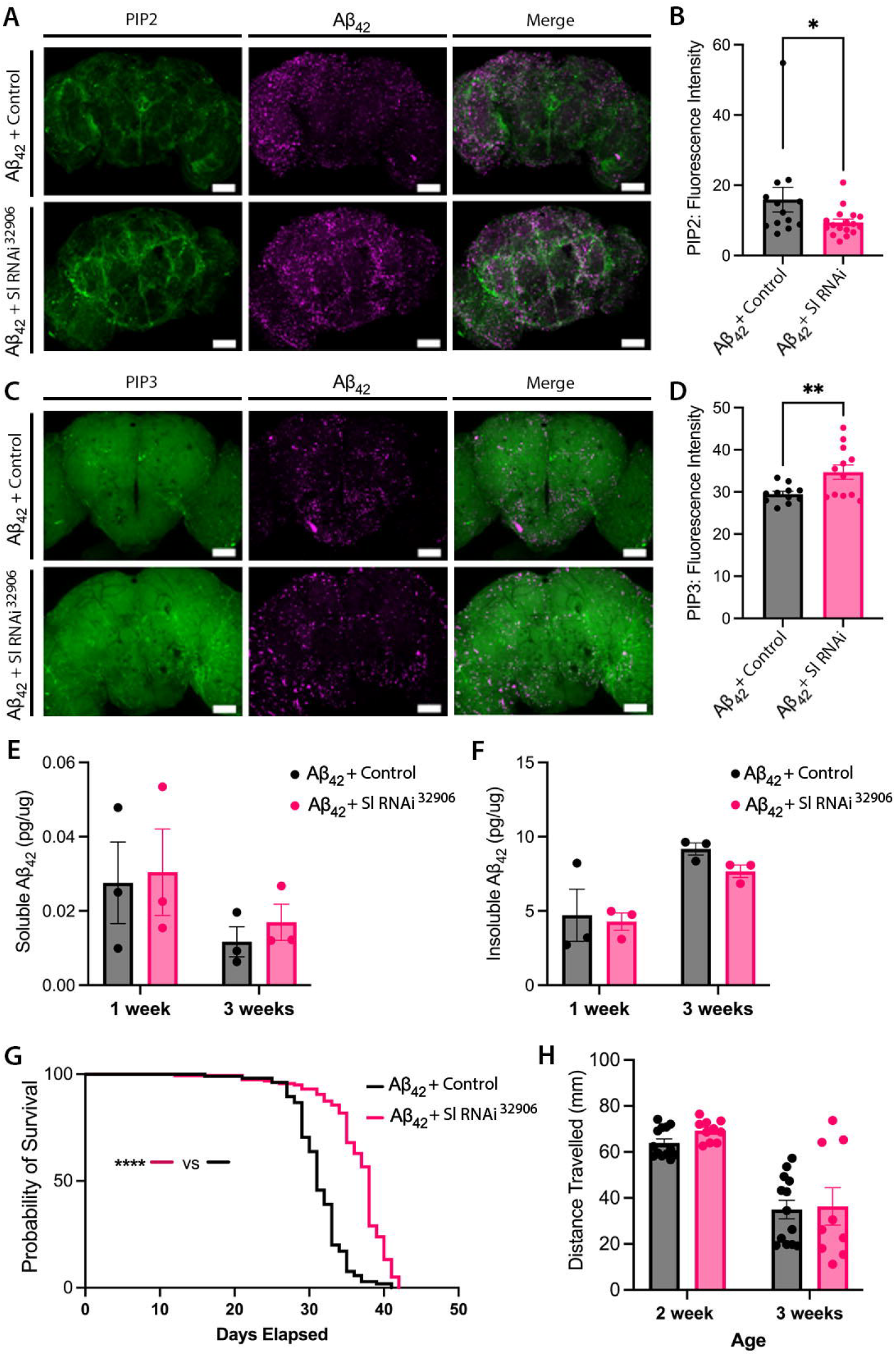
Small wing knock down modifies phosphoinositide homeostasis and survival in a *Drosophila* Aβ42 overexpression model. **A)** Representative images showing glial restricted (repo-Gal4) expression of secreted Aβ_42_ ^Arctic^ (Magenta) and PIP2 fluorescent reporter (PLCd-PH::GFP, Green) in 7-dpe adult fly brains. n=12-18 brains per group. B) Quantification of PIP2 fluorescent reporter signal intensity revealed a modest downregulation in flies expressing RNAi targeting *sl*. C) Representative images showing glial restricted expression of Aβ_42_ ^Arctic^ (Magenta) and PIP3 fluorescent reporter (*GRP1-PH::GFP*, Green) in 7dpe adult fly brains. D) Quantification of PIP3 fluorescent reporter signal intensity revealed an increase in flies expressing RNAi targeting *sl*. E) Levels of Aβ_42_ from soluble and F) insoluble extracts of fly heads were measured using Meso Scale Discovery quantitative immunoassay, detecting no change in amyloid levels between sl RNAi and GFP-Valium10 control flies. Three independent biological replicates of 40 adult heads per genotype. G) Lifespan associated with glial expression of Aβ_42_ ^Arctic^ is extended by *sl* knockdown relative to controls (24 hours mated females, n=105-159). Imaging was analysed by One-way ANOVA with Dunnett’s multiple comparison test and Lifespan by Log Rank, Mantel-cox test Graphs show mean±SEM, *p<0.05, **p<0.01, ****p<0.0001. Scale bar: 50 µm.

To assess the impact of *sl* knockdown on Aβ_42_-relevant phenotypes of whole organism fitness, we next measured longevity and locomotion function. Despite not impacting on amyloid accumulation, we found that *sl* knockdown improved the survival of flies expressing Aβ_42_, with an increased median survivorship of ∼10 days (p<0.0001), (Fig 5G). Effects seem to be amyloid specific as we previously show *sl* knockdown in glia has no effect to longevity in wildtype flies (Fig. 1F). Improved viability associated with glial downregulation of *sl* was not however accompanied by significant changes in neuronal function. Age-associated decline in motor performance in a negative geotaxis assay remained consistent between control and *sl* RNAi expressing flies regardless of age tested (Fig 5H). Taken together, these data indicate that *sl* mediated PIP2 metabolism in glia contributes to longevity of flies expressing amyloid beta, however does so independently of direct modification of amyloid burden.

Given the association of *PLCG2* coding variants with late-onset AD, we generated transgenic flies expressing the common human *hPLCG2-P522* variant, the protective *hPLCG2-R522* variant, and *sl*. Increased glial expression of either human *PLCG2* variant or *sl* in a wildtype background had no detrimental impact on longevity, while both human variants maintained normal motor function with age (Supplementary material, Fig. 3). In an Aβ_42_ background, glial overexpression of *sl* or the common *hPLCG2-P522* variant significantly reduced survival, whereas the protective *hPLCG2-R522* variant preserved lifespan comparable to controls. Motor deficits induced by Aβ_42_ were not significantly altered by expression of either human *PLCG2* variant, suggesting that the protective R522 variant selectively modulates survival-related pathways in response to amyloid toxicity.

## Discussion

Efficient clearance of degenerating neuronal material is a fundamental function of glia required for maintenance of neural homeostasis and recovery following injury. Here, we identify phosphoinositide turnover through the *Drosophila* PLCγ orthologue *small wing* (*sl*) as a key regulator of glial engulfment downstream of the engulfment receptor Draper. We show that glial depletion of *sl* profoundly delays removal of degenerating axons, disrupts injury-induced phosphoinositide remodelling, attenuates Draper upregulation and alters calcium signalling responses following neuronal injury. Together, these findings support a model in which PLCγ-dependent phosphoinositide dynamics coordinate membrane and signalling changes required for efficient glial phagocytosis.

A growing body of human genetic evidence implicates phosphoinositide metabolism in neurodegenerative disease. Variants in *PLCG2* and *INPP5D* are associated with altered risk of late-onset Alzheimer’s disease, placing phosphoinositide signalling among the major microglial pathways linked to disease susceptibility (Bellenguez et al., 2022; Sims et al., 2017). PLCG2 is highly enriched in microglia and functions downstream of immune receptors including TREM2, where it contributes to intracellular calcium release, lipid signalling and phagocytic responses (Andreone et al., 2020; Tsai et al., 2023). Previous mammalian studies have demonstrated that PLCG2 regulates microglial survival, migration and amyloid-associated responses, including plaque compaction and phagocytosis (Maguire et al., 2021; Takalo et al., 2020). However, whether PLCγ signalling directly regulates engulfment of degenerating neuronal material *in vivo* has remained unclear. Consistent with a role in maintaining glial clearance capacity, loss of *sl* reduced the expression of genes associated with host defence and phagocytic responses, indicating that PLCγ activity influences both the cellular and transcriptional programmes that support engulfment. Our findings provide evidence that this pathway plays a conserved role in glial engulfment following axonal injury.

The engulfment defects observed following glial *sl* depletion resemble phenotypes associated with disruption of Draper signalling, the principal phagocytic receptor controlling glial clearance of degenerating axons in *Drosophila*. Draper functions analogously to mammalian MEGF10 and activates downstream signalling pathways involving Src42A, Shark and Rac1 to drive cytoskeletal rearrangement and phagocytic engulfment (Doherty et al., 2014b; Logan et al., 2012; MacDonald et al., 2006; Ziegenfuss et al., 2008). Consistent with this established pathway, we found that loss of either *Draper* or *sl* abolished the normal injury-induced increase in membrane-associated PIP3. These findings position phosphoinositide remodelling downstream of engulfment receptor activation and suggest that PLCγ activity is required to coordinate the membrane lipid transitions necessary for phagosome formation and engulfment progression. Previous work has demonstrated that phosphoinositides regulate multiple stages of phagocytosis, including receptor clustering, actin remodelling and phagosome maturation, through highly localised changes in membrane identity (Gillooly et al., 2001; Levin et al., 2015). Our data extend these concepts to glial engulfment *in vivo* and identify PLCγ-dependent phosphoinositide turnover as an important component of Draper-mediated injury responses.

One notable finding was that *sl* depletion elevated basal PIP2 levels while preventing injury-induced accumulation of PIP3. PLC enzymes hydrolyse PI(4,5)P2 to generate DAG and IP3, thereby linking membrane phosphoinositide metabolism to calcium signalling and downstream activation pathways. Previous work identified an interaction between a *Drosophila* PI3K adaptor protein and PLCγ, suggesting that PI3K signalling may contribute to PLCγ localisation and activation (Weinkove et al., 1997). Recruitment of PLCγ to membrane phosphoinositide signalling domains has also been demonstrated in mammalian systems (Falasca et al., 1998). Elevated basal PIP2 in *sl*-deficient glia is consistent with reduced phospholipase activity. Importantly, simultaneous knockdown of *pten*, which converts PIP3 back to PIP2, rescued delayed engulfment caused by *sl* depletion. This genetic interaction strongly supports disrupted PIP2/PIP3 balance as a mechanistic contributor to impaired engulfment. PI3K-generated PIP3 is known to recruit proteins involved in actin polymerisation and membrane extension during phagocytosis, while PTEN spatially restricts these signalling domains (Desale & Chinnathambi, 2021; Lennartz, 1999; Mondal et al., 2011; Schlam et al., 2015). Our findings suggest that insufficient PIP3 accumulation in *sl*-deficient glia limits effective engulfment and that restoring phosphoinositide balance can compensate for reduced PLCγ activity.

In addition to altered phosphoinositide dynamics, *sl* depletion prevented normal injury-induced upregulation of Draper and disrupted calcium responses following axotomy. Calcium signalling is a critical downstream consequence of PLC activation through IP3-mediated release of ER calcium stores and contributes to cytoskeletal remodelling, vesicle trafficking and phagosome maturation. Interestingly, glial *sl* knockdown increased baseline calcium levels yet abolished further calcium elevation following injury, suggesting dysregulated calcium homeostasis rather than simple loss of signalling capacity. Similar alterations in calcium signalling downstream of TREM2–PLCG2 pathways have been observed in mammalian microglia (Andreone et al., 2020). Together, these findings support the idea that PLCγ signalling acts as a central coordinator coupling engulfment receptor activation to membrane phosphoinositide turnover and calcium-dependent phagocytic machinery.

Finally, we demonstrate that manipulating glial phosphoinositide signalling modifies organismal outcomes in a fly model of amyloid pathology. Reduced glial *sl* expression improved survival independently of detectable changes in soluble or insoluble Aβ42 levels, suggesting that altered glial signalling states can influence disease-relevant phenotypes without directly modifying amyloid burden. Moreover, glial expression of human *PLCG2* variants differentially altered longevity in Aβ42-expressing flies, with the AD-protective R522 variant preserving lifespan relative to the common P522 allele. The R522 variant has been proposed to function as a hypermorphic allele that enhances PLCG2 activity and microglial responses (Maguire et al., 2021; Takalo et al., 2020). Our data support the idea that subtle modulation of PLCγ signalling influences glial functional states relevant to neurodegeneration. Importantly, both reduced and enhanced PLCγ activity may produce context-dependent effects, highlighting the importance of balanced phosphoinositide signalling for glial homeostasis.

In summary, our findings identify PLCγ-dependent phosphoinositide turnover as a conserved regulator of Draper-mediated glial engulfment. We propose that neuronal injury activates Draper signalling to drive local phosphoinositide remodelling, calcium signalling and receptor upregulation through *sl*/PLCγ activity, thereby enabling efficient clearance of degenerating neuronal material. Given the strong genetic association between *PLCG2* and

Alzheimer’s disease, these findings provide mechanistic insight into how altered phosphoinositide signalling may influence glial states and neurodegenerative disease progression.

## Author Contributions

FS, EM, GS and OP conceived ideas, wrote the paper, and did experiments; HC, DM, BM and LA, TS did experiments; FS and EM were supervised by GAS and OP.

## Funding

This work was funded by the MRC Momentum Award (MC_PC_16030/1 to GAS), Leverhulme Trust project grant (RPG-2020-369 to GAS), MRC Momentum Award (MC_PC_16030/2 to OP), MRC project grant (MR/W004879/1 to OP) and Wellcome Trust Integrative Neuroscience PhD studentship (108891/B/15/Z to HC & GS).

## Conflicts of Interest

The authors declare no conflicts of interest.

## Data Availability Statement

Large data sets and analysis codes have been deposited open-source platforms as outlined in the method section. Additional datasets and materials generated during the current study are available from the corresponding author upon reasonable request.

## Notes

https://github.com/Fly-Cardiff/Fly-RING-Analysis

https://github.com/Fly-Cardiff/RNA_Seq

https://www.ebi.ac.uk/biostudies/ArrayExpress/studies/E-MTAB-14080?query=E-MTAB-14080

### Competing Interest Statement

The authors have declared no competing interest.

