## Supplementary material for "Phosphoinositide turnover through PLCγ regulates Draper-dependent engulfment in glia"

Hadyn Ellis Building

Cardiff University

Maindy Road

Cardiff

Wales, UK, CF24 4HQ

**Supplementary Methods**

**Quantitative PCR**

*Sl* RNAi and control lineswere crossed to *tubulin*-GAL4and raised at 29^o^C. 12-24 hrs post eclosion, flies were snap-frozen in liquid nitrogen. RNA was isolated from whole flies homogenised in TRIzol reagent (Invitrogen, 15596026), according to manufacturer’s instructions. Genomic DNA contamination was removed by TURBO DNase kit (Invitrogen, AM1907) and cDNA was synthesised using the QuantiTect Reverse Transcription kit (Thermo Scientific, K0221) according to manufacturer’s instructions. cDNA was amplified using Maxima SYBR Green/ROX master mix (Invitrogen) and the following primers at 300nM final concentration; *sl* forward*: ACGGCCACACCATTACCTCC, sl* reverse: *CGGGGATATGCTGCTTACGC*, *rp49* forward:*AGCATACAGGCCCAAGATCG, rp49* reverse: *TGTTGTCGATACCCTTGGGC.* The QuantStudio 7 thermocycler was used with the following conditions: Initial denaturation (95^o^C, 600 s), denaturation (95^o^C, 15 s), annealing (60^o^C, 30 s), extension (72^o^C, 30 s) (40 cycles), melt curve (60-95^o^C). Melt curve analysis was performed to ensure single-product amplification. Raw amplification data was processed using the qpcR package in R Studio and statistical significance was calculated using a permutations approach (Ritz & Spiess, 2008).

**Wing Size Quantification**

*Sl* RNAi lines (BDSC 32906 and VDRC 108593) were crossed to the ubiquitous *Tub-Gal4* promoter and reared at 25°C. Male and female flies from the F1 progeny were selected and aged for 5 days at 29°C prior to wing dissection. 10 wings per genotype were mounted on to a glass slide with coverslip. Digital images of the wings were taken on a Zeiss Stemi 508 microscope connected to a 5.0 Mega Pixel camera (AxioCam ERc 5s, Carl Zeiss) and analysed using the Fiji software (ImageJ). A region of interest was drawn around the L4, L5 and posterior cross vein perimeter and area measured.

**Molecular Cloning and Fly Stock generation**

To generate site directed insertion fly lines, cDNA encoding the *Drosophila* *sl* gene (DGRC, GOLD RE62235) and human *PLCG2* gene (DGRC, HSCD00506018) were subcloned into the *5xUAS-pJFRC5* vector (Gifted from Gerald Ruben, Addgene, plasmid 26218). *sl* and *PLCG2* cDNA were amplified by PCR using primer sequences: *sl* forward TCAGCGGCCGCACAACCAAAATGAGCTGCTTTAGTGCGAT, *sl* reverse GTATCTAGACTACGGTGCGGTAACATTTG, *PLCG2* forward TCAGCGGCCGCACAACCAAAATGTCCACCACGGTCAATGT, *PLCG2* reverse GTATCTAGACTACGGTGCGGTAACATTTG that added NotI and XBaI restriction sites to the 5’ and 3’ ends. Amplified PCR product was isolated from agarose gels according to manufacturer's instructions (Thermo Fisher, K220001). 1µg of *5xUAS-pJFRC5* vector, *sl* and *PLCG2* DNA were digested with *Not1-HF* and *XBaI* restriction enzymes and ligated using T4 DNA ligase (New England Biolabs, M0202S), with addition of alkaline phosphatase to prevent re-ligation of the vector backbone. DH5a cells (Thermo Fisher, 18263021) were transformed with ligation product and individual colonies were picked from LB + Ampicillin resistance plates. Plasmid DNA from positive colonies was isolated using the QIAprep Spin Miniprep kit (Qiagen, 27106) and sequence verified (Genewiz). For site-directed mutagenesis to generate *PLCG2-R552*, 4µg of plasmid containing wildtype *PLCG2-P552* DNA was provided to commercial vendor Genscript (Netherlands, Leiden). Site-directed mutagenesis introduced a C to G base substitution at position 1564bp of the *PLCG2* insert, resulting in a Proline to Arginine conversion.

***Drosophila* embryo injection for transgenic production**

Embryo injection and site directed integration of verified *sl* and *PLCG2* variant plasmids was conducted by a commercial vendor (BestGene, USA). *Drosophila melanogaster* embryos from BDSC 36304 & 8622 stocks, harbouring attP40 and attP2 landing sites, respectively, were Injected constructs were integrated using PhiC31 integrase-mediated site-specific recombination. An empty JFRC vector control line was generated in parallel and used as a transgenic insertion control throughout the study. Successful transformants were identified, back-crossed into a w1118 background, and balanced prior to experimental use.

**Supplementary Figure 1**

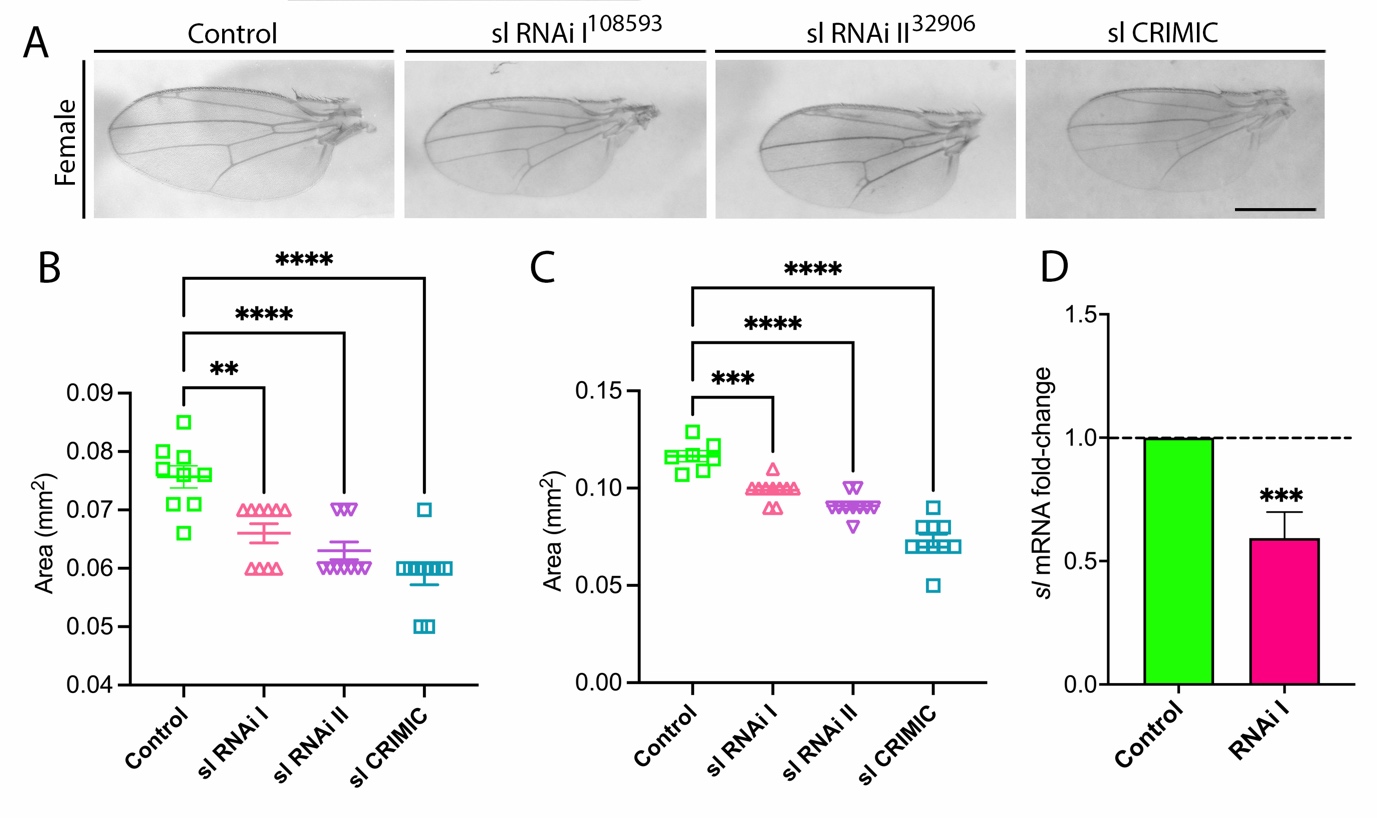

**Supplementary Figure 1. Verification of RNAi mediated knockdown of small wing.**

A) Ubiquitous expression (tubulin-Gal4) of two independent sI targeting RNAi, or ablation of sl expression by a CRIMIC null mutant, give rise to a developmental phenotype of anatomically smaller wings. Quantification of wing segment contained within L4, L5 and posterior cross vein perimeter demonstrating significantly smaller wing development in (B) female and (C) male adult flies. D) Quantitative PCR analysis of ubiquitous (Tubulin-Gal4) sl RNAi 1 expression demonstrates significant knock down in the adult fly head. Statistical analysis was achieved through a one-way ANOVA with Dunnett’s multiple comparison test. Graphs were plotted with mean±SEM, ***p<0.001 and ****p<0.0001, n=7-10 wings per group. Scale bars = 0.6 mm.

**Supplementary Figure 2.**

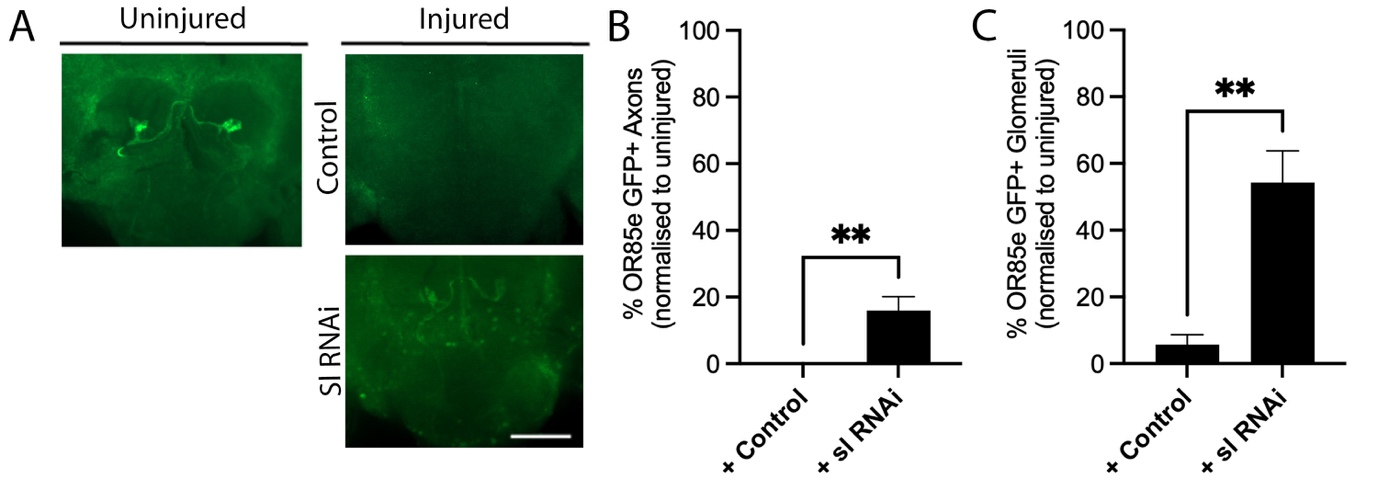

**Supplemental Figure 2. Small wing contributes to glial engulfment of neuronal debris.**

A) GFP-labelled maxillary olfactory receptor neurons following axotomy in control (attP2) or following glail specific knockdown of sl flies at 3 days after injury. Data derived from second independent RNAi (II). B) Analysis of percentage of antennal lobes containing visible GFP+ axons derived from severed OR85e maxillary ORN neurons and C) GFP+ glomeruli detected significantly delayed clearance in *sl* RNAi expressing flies. Data was analysed using T-test. Graphs show mean±SEM, **p<0.01, n=7-10 brains per group. Scale bars = 50μm.

**Supplementary Figure 3**

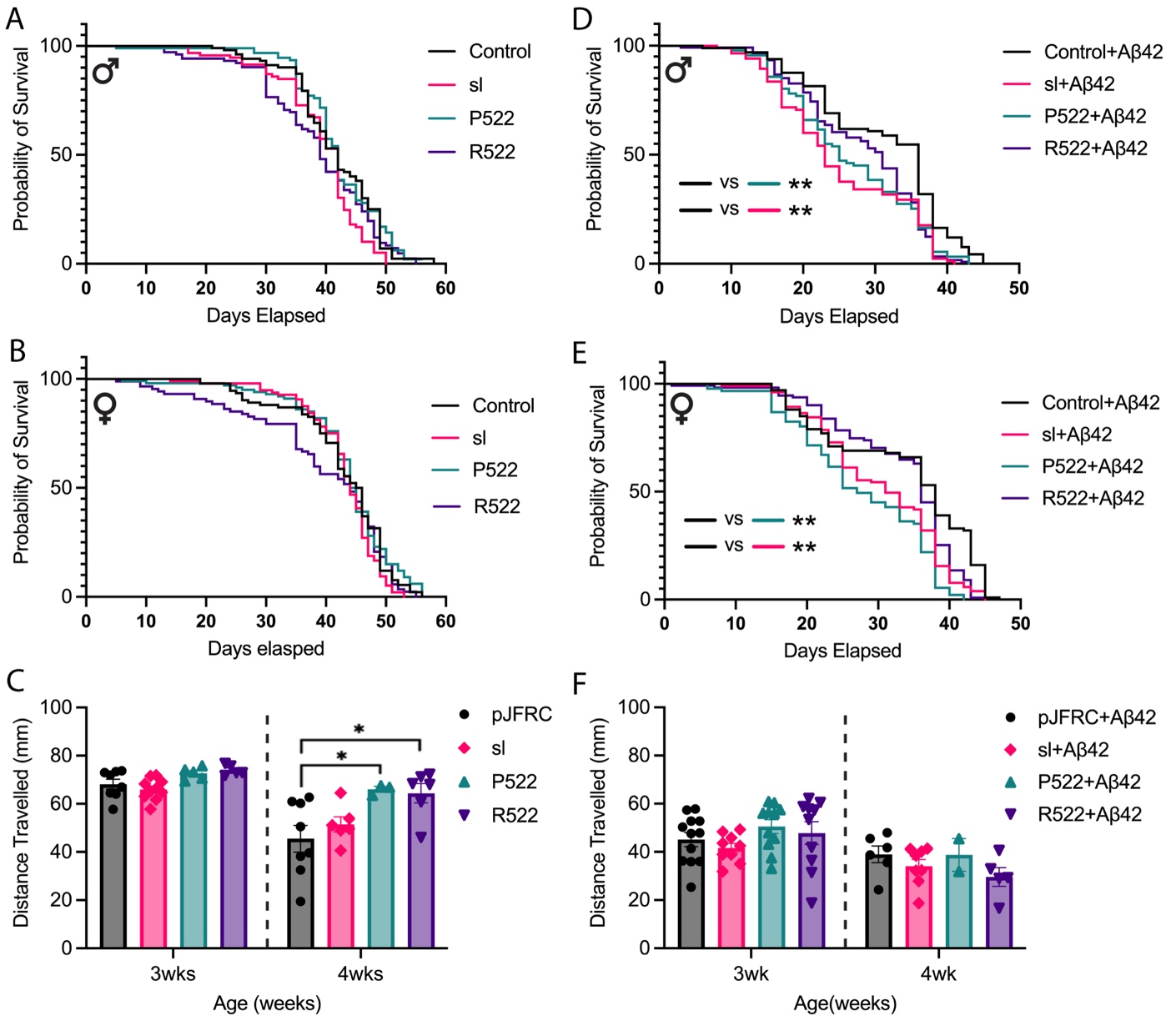

**Supplementary Figure 3. Roles for glial sl and PLCG2 in survivorship and locomotor function**

No differences in survival of A) male and B) female (24 hours mated) flies with glial restricted expression of expression of sl, human PLCG2 variants (common P522 and rare R522) or a control empty pJFRC5 vector. (n=182-265, Log-rank, Mantel-Cox test). C) Average distance travelled (mm) in 10 seconds post negative geotaxis recorded for both female and male flies at 3-week timepoints (n=5-10 vials, one-way ANOVA with Sidak’s multiple comparison test) In both D) male and E) female flies, glial co-expression of Aβ_42_^Arctic^ and either *Drosophila* sl or human PLCG2 P522 reduced lifespan, with PLCG2-R552 maintaining control lifespan. F) Neither glial expression of sl or human PLCG2 variants impacted on RING motor behaviour performance in Aβ_42_^Arctic^ transgenic flies. Graphs show mean±SEM, *p<0.05, **p<0.01
